# Gender stereotype and the scientific career of women: Evidence from biomedical research centers

**DOI:** 10.1101/2020.10.29.360560

**Authors:** Jose G. Montalvo, Daniele Alimonti, Sonja Reiland, Isabelle Vernos

**Affiliations:** Universitat Pompeu Fabra, Department of Economics and Business, Universitat Pompeu Fabra, Ramon Trias Fargas 25-27, Barcelona, 08005, Spain; Barcelona Graduate School of Economics, Ramon Trias Fargas 25-27, Barcelona, 08005, Spain; Universitat Pompeu Fabra, Department of Economics and Business, Universitat Pompeu Fabra, Ramon Trias Fargas 25-27, Barcelona, 08005, Spain; Institute of Political Economy and Governance, Ramon Trias Fargas 25-27, Barcelona, 08005, Spain; Center for Genomic Regulation, Dr. Aiguader 88, Barcelona 08003, Spain; The Barcelona Institute of Science and Technology, Carrer del Comte d’Urgell 187, Barcelona 08036, Spain; Center for Genomic Regulation, Dr. Aiguader 88, Barcelona 08003, Spain; Universitat Pompeu Fabra, Barcelona, Spain; Institució Catalana de Recerca i Estudis Avançats, Pg. Lluis Companys 23, Barcelona 08010, Spain

## Abstract

Women are underrepresented in the top ranks of the scientific career, including the biomedical disciplines. This is not generally the result of explicit and easily recognizable gender biases but the outcome of decisions with many components of unconscious nature that are difficult to assess. Evidence suggests that implicit gender stereotypes influence perceptions as well as decisions. To explore these potential reasons of women’s underrepresentation in life sciences we analyzed the outcome of gender-science and gender-career Implicit Association Tests (IAT) taken by 2,589 scientists working in high profile biomedical research centers. We found that male-science association is less pronounced among researchers than in the general population (34% below the level of the general population). However, this difference is mostly explained by the low level of the IAT score among female researchers. Despite the highly meritocratic view of the academic career, male scientists have a high level of male-science association (261% the level among women scientists), similar to the general population.

## 1 Introduction

Dozens of indicators show that women are underrepresented in public life and managerial positions in private companies. Women are also underrepresented in the academic community: while there is an increasing number of Ph.D. degrees awarded to women, the proportion of women decreases from the postdoctoral level to assistant professors, and ever further, to full professor positions. For instance women obtain 45% of doctorates in science, mathematics and engineering but they only represent 30% of the senior faculty (National Science Foundation (US), 2016). This phenomenon, pervasive in other disciplines such as biology (National Science Foundation (US), 2018), is sometimes referred to as a “leaky pipeline” although this view is not shared by all the literature (Kang & Kaplan, 2019). Women account for only 33% of the researchers of the EU, 24% of the top-level researchers, and 27% of board leaders in scientific boards (European Commission, 2019).

Many indicators suggest the existence of a certain degree of gender bias in science at many different levels including publishing, hiring, funding, and earnings. Female scientists get less citations for papers of similar quality (Caplar, Tacchella, & Birrer, 2017; Ghiasi, Larivière, & Sugimoto, 2015), even if they are in dominant author positions (Larivière, Ni, Gingras, Cronin, & Sugimoto, 2013); women are underrepresented in the peer-review process (Helmer, Schottdorf, Neef, & Battaglia, 2017; Wennerds & Wold, 1997); they are less often described as “brilliant” and above others than their male counterparts in recommendation letters for postdoctoral fellowships (Dutt, Pfaff, Bernstein, Dillard, & Block, 2016); they are hired as independent fellows in bio-medical research at a lower rate than males (Sheltzer, 2018) and elite male faculty trained significantly fewer women than other female faculty members (Sheltzer & Smith, 2014); women also have less success in grant applications than men if research proposals are judged on the strength of their CV (Ley & Hamilton, 2008). There is also evidence that female scientists receive a lower salary than their male counterpart (Shen, 2013). Recent studies, using fictional CVs, have provided strong evidence for gender bias playing a major role in these inequalities(Moss-Racusin, Dovidio, Brescoll, Graham, & Handelsman, 2012; Reuben, Sapienza, & Zingales, 2014). And there is evidence that this situation has not been improving significantly over time (Helmer et al., 2017).

Similar patterns of general underrepresentation of women in academic positions in general are also observed in the biomedical sciences. Indeed, women get the majority of the doctoral degrees in fields related with biology but they only represent one third of the assistant professors in these disciplines (National Science Foundation (US), 2018). These patterns are also very similar in different geographical areas. For instance, in the European Union, where we focused our study, women researchers have a success rate in funding applications which is 3 percentage points smaller than men (European Commission, 2019).

Recent research shows (Lerchenmueller & Sorenson, 2018) that women become principal investigators (PI) at a 20 % lower rate than men in life sciences, and discuss the so-called “productivity paradox”. Lerchenmueller & Sorenson, 2018 show that differences in the publication record and citations per article can explain about 60 % of the probability of receiving an NIH R01 grants while the differential return on the published publications and, in particular, women receiving less credit for their citations, could explain most of the remaining part.

But, why do women publish less than men? Why do women get less citations than men? Several competing theories have tried to explain this persistent phenomenon, such as putative differences in innate aptitudes among men and women at the high end of the scientific aptitude distribution, which is currently loosing support through increasing evidence against the innate attitude theories (Hyde & Mertz, 2009; Hyde, Lindberg, Linn, Ellis, & Williams, 2008); different career preferences, due to a number of considerations such as for instance the willingness to work long hours or the disposition for service jobs, which reduces the time available for research and increases the likelihood to decline participation in peer-evaluations (Helmer et al., 2017); and gender bias / discrimination exemplified by negative stereotypes with respect to scientific ability and talent. There is in fact mounting evidence for these negative stereotypes associated with women, as we show in the next section. Results from a survey of US academics showed that female are underrepresented in disciplines where professors believe that innate talent is the basic trait for success, and women are stereotyped as not displaying such a talent (Hyde & Mertz, 2009). But, do scientists exhibit a higher degree of negative stereotypes associated with women than the general population?

## 2 The unconscious nature of gender bias

In many instances negative stereotypes are described as an “unconscious bias” (Mervis, 2012; Reuben, 2014). In general, women’s underrepresentation in science is not the result of explicit and easily recognizable gender biases but the outcome of decisions with many components of unconscious nature that are difficult to assess and address (Helmer et al., 2017). Evidence suggests that gender stereotypes influence perceptions and performance as well as decisions. Recent theories in social cognition recognize that implicit measures of stereotyping are more automatic, and less conscious, than the explicit measures derived from self-reported assessments obtained using surveys. Implicit measures do not require introspection on the part of the subject and, therefore, generate less skepticism than self-evaluations.

Since the biases that play a role in the underrepresentation of women in the scientific community have mostly an unconscious nature there is a need for an instrument that can capture this implicit bias. One very successful instrument to measure the association of stereotypes to particular groups of the population is the Implicit Association Tests (IAT). The IAT measures, using the response latencies and accuracy, the strength of automatic association of combined categories (e.g. male vs. female) and attributes stimuli (e.g. good vs. bad). Since the original publication of the IAT in 1998 (Greenwald, McGhee, & Schwartz, 1998), many studies have established construct validity (Nosek, Greenwald, & Banaji, 2005; K. A. Lane, Banaji, Nosek, & Greenwald, 2007) and dissociation between implicit and explicit measures (Hofmann, Gawronski, Gschwendner, Le, & Schmitt, 2005)^1^.

The evidence of implicit gender stereotypes related to science and career has accumulated over time. Research has found that both, men and women, are more likely to associate male and science, and that men are more likely to pursue science careers whereas women are more likely to pursue careers in humanities (K. Lane, Goh, & Driver-Linn, 2012). Country differences in gender-science stereotype can predict gender differences in science and math achievement (Nosek et al., 2009). Women implicitly identify with their gender group, and associate female with liberal arts, which in turn promotes their preference for liberal arts (Nosek, Banaji, & Greenwald, 2002). Although female engineering students held weaker implicit gender-math and gender-reasoning stereotypes than female students in humanities and male students in engineering or humanities (Smeding, 2012), the hypothesis that women working in scientific disciplines have weakened the science-is-male stereotype is not supported by the available evidence (Miller, Eagly, & Linn, 2015; Smyth & Nosek, 2015). In particular, biological science majors report a weaker explicit male-science association than other majors, but the same level of implicit association (Smyth & Nosek, 2015). Moreover, stronger gender-science stereotypes were found to be associated with weaker science identification and, in turn, weaker science career aspirations (Cundiff, Vescio, Loken, & Lo, 2013).

## 3 Basic results

In this study we address two basic questions: Are scientists working at top biomedical research centers also subjected to the influence of gender stereotypes with respect to science and scientific career? Do active researchers show more or less gender-science associations than the general population? We used the gender-science and gender-career Implicit Association Tests (IAT) to elicit the implicit attitudes in a large sample of active scientists. The sample contains more than 2,500 researchers working in 13 high profile life science research centers located in European countries and belonging to the EuLife alliance. These centers collaborate in the Leading Innovative Measures to Reach Gender Balance in Research (LIBRA) project through which this study was implemented. Employees at participating centers^2^ were asked to take the two computer-based IAT tests anonymously using the online platform and procedures of the Implicit Project.

To assess the work environment, and the possibility of interacting sociocultural factors associated with the gender composition of the workforce, we also obtained general information on gender and professional indicators for each of the collaborating centers (percentage of women among researchers, proportion of women among IPs, etc.). The determinants of the scores were also compared with the factors explaining them in the general population. To characterize this group we used the results from individuals who took the tests for the Project Implicit. The results are robust to restricting the sample to Europeans over 26 years old, which is a subsample closer to the basic demographic characteristics of our scientists than the general sample. It should be noted that, since implicit attitudes could potentially change over time (Charlesworth & Banaji, 2019) we only considered the tests taken during the period in which the scientists took the IAT (2015-17). For this period the sample for the general population includes 320,783 individuals for the gender-science IAT and 572,262 for the gender-career IAT.

The paper contains two important novelties with respect to the previous literature. First, we present the results of Implicit Association Tests taken by a large group of researchers working in elite biomedical research centers in Europe. It includes researchers at all the stages of their professional career which allows, given the large sample size, to analyze implicit association for different professional positions. We also analyze the individual and center determinants of their implicit association scores. Our sample is restricted to researchers in life sciences since recent research shows that there are large difference in implicit bias by academic major (Smyth & Nosek, 2015). This strategy avoids pulling together researchers from different research fields, and potentially very different implicit associations. As far as we know this is the first time that the Implicit Association Test has been taken by such a large number of active researchers in top research centers of life sciences. Since gender bias can be described in many situations as an unconscious effect we believe our analysis is very relevant for studying the persistent issue of the low proportion of women in top scientific positions. Recent research has shown that, in evaluation committees, the stronger the implicit stereotype, the less often women are promoted (Régner, Thinus-Blanc, Netter, Schmader, & Huguet, 2019). Second, we compare the IAT scores for the gender-science and gender-career associations of researchers with those obtained by the general population. The objective is not only to compare the difference in the raw scores but also to examine the influence of different factors (gender, age, etc.) on the implicit association between gender and science (and gender and career).

First of all, we looked into the gender composition of the centers by professional positions. Overall the proportion of women in these research center ranges from 42.1% to 66.0%. The population of researchers consists of 52.5% of women and 47.5% of men. However, Figure 1 shows that the proportion of women decreases with rank. It is the highest for the administrative personnel (70%) and research staff (61.6%) and it goes down to 19.3% for group leaders. If we consider the proportion of women in top roles (group leaders and heads of units), they account only for 21.1% of the positions. This figure is very close to the average proportion of women reported for top level researchers in the European Union (European Commission, 2019). Therefore, the centers included in the study are quite representative of research activities in the EU.

**Figure 1:**
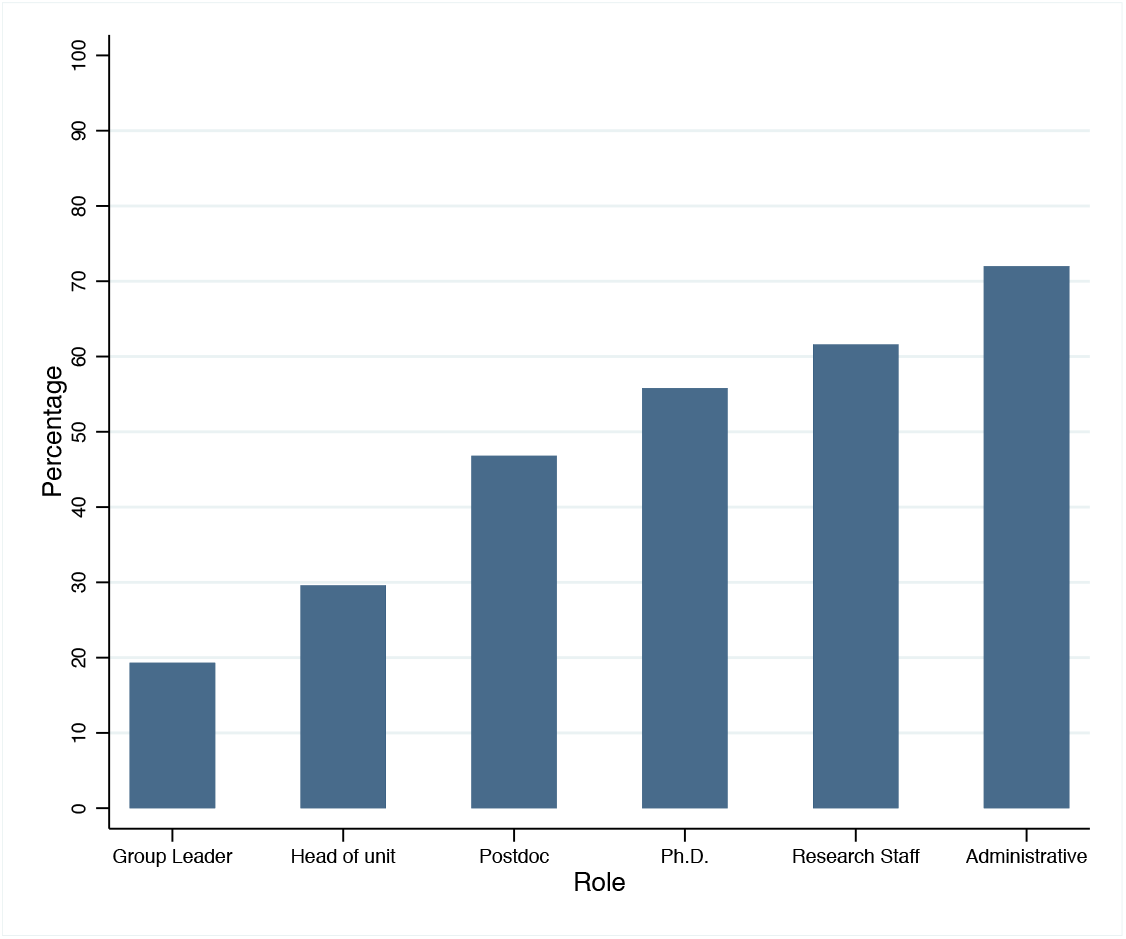
Percentage of women by professional category.

As shown in the literature, the available evidence points towards implicit gender bias in many population groups, including college graduates and university students. The novelty of our paper is to extend these results to active research scientists working in very competitive research centers, where merit should be the main criterion for selection and promotion, and to compare the results with the general population. In principle, being socialized in this kind of environment should provide scientists with some protection against gender-science associations.

Gender-science and a gender-career Implicit Association Tests (IAT) were completed by the subjects working at the research centers, and their performance in the tasks recorded. The main statistic used to analyzed the IAT is the D-score. To compute these scores we use the score algorithm that has been shown to have the best performance (Greenwald, Nosek, & Banaji, 2003). In particular, data from all blocks of the test were used. Individual latencies greater than 10,000 ms were eliminated as well as subjects for whom more than 10% of trials have latency less than 300 ms. Using all the trials, we computed the mean of each block as well as one pooled standard deviation for all trials in the first and third block and another for the second and fourth block. Response times when the answer was an error was replaced by the mean of the block plus a penalty of 600 ms. Then, we average the values of each block before computing the differences between the third and the first block and the last and the second block. The reaction time differences between stereotype consistent and inconsistent blocks were divided by the standard deviations of the latency and the two quotients were averaged out to compute the D-score. The interpretation of the score is simple: positive scores implied male-science associations while negative scores implied female-science association.

The D-scores of the gender-science IAT show evidence of gender stereotyping both among biomedical researchers (Project LIBRA) and general population (Project Implicit). We consider the subsample of LIBRA that includes only scientist (excluding the administrative staff working at the centers)^3^. Table 1 shows that the score for the sample of scientists is 0.23, 95% CI[.21, .24], and for the participants in the Project Implicit is 0.312, 95% CI[.311, .313]. The internal consistency between blocks shows that the measurements are robust, with correlations above the standard figures found in gender-science IAT (Greenwald et al., 2003). The order effect, or sensitivity of the score to the order of the blocks in the test, is also smaller than the usual value in the gender-science IAT among the general population (Greenwald et al., 2003). These tests show that the measurements of the IAT are robust.

**Table 1:**
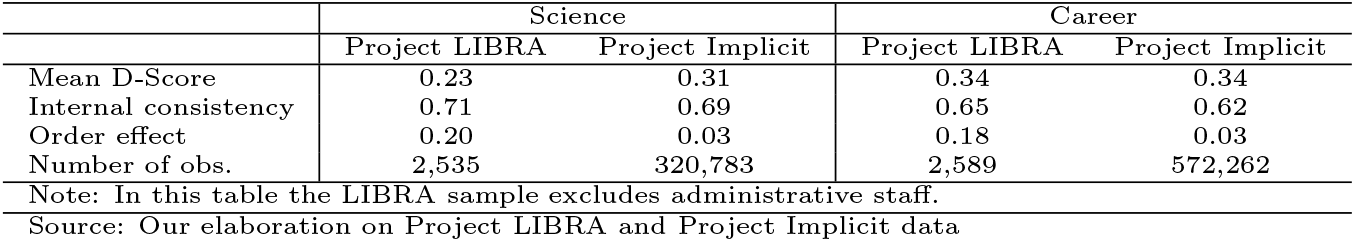
Comparison between Project LIBRA and Project Implicit.

Are workers at research center more or less gender biased than the general population? To answer this question we compared the basic results of the biomedical researchers working in top EU research centers with the results from the general population. Given the large size of the general population sample we report size effects, mostly Cohen’s ds. Cohen’s *d* measures the effect-size parameter of interest in the scaled difference between means

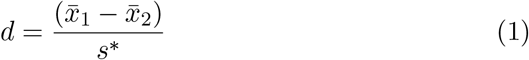

where

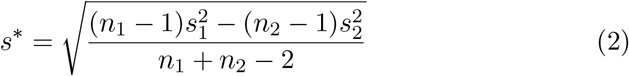

Since Cohen’s d is biased there is an unbiased version, Hedges’ g, which takes the form *g=d.c(m)* where *m* = *n*_1_ + *n*_2_ − 2

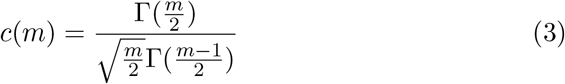

Table 1 shows that the gender-science IAT score of scientists is lower than the score of the general population, while for gender and career the scores are identical. Cohen’s d indicates that the average score of the IAT for science differs between the sample of the general population (project Implicit) and the sample of researchers (project LIBRA) by 0.18 standard deviations with a 95% CI[0.14, 0.22]. Table 2 breaks the results by gender^4^. It shows that most of the lower implicit association of gender and science among scientists is actually due to the lower implicit association of women in research centers with respect to women in the general population, while male researchers have a similar score to the general male population^5^. In particular, Cohen’s d points out that the difference in the science IAT of males comparing the general population and the researchers is 0.05 with a 95% CI[−0.01, 0.12] while Cohen’s d for the comparison between females in the general population and in the sample of researchers is 0.26 with a 95% CI[0.18, 0.34]. The difference between men and women at the research centers in the science IAT is 0.42 standard deviations with a 95% CI[0.50, 0.34]. In this set-up the previous Cohen’s ds are almost identical to their corresponding Hedges’ gs.

**Table 2:** Comparison by Gender.

|  | Project LIBRA |  | Project Implicit |  |
| --- | --- | --- | --- | --- |
|  | Female | Male | Female | Male |
| Mean D-score: Science | 0.13 | 0.34 | 0.28 | 0.36 |
| Mean D-score: Career | 0.36 | 0.27 | 0.38 | 0.29 |

Note: In this table the LIBRA sample excludes administrative staff.
Our elaboration on Project LIBRA and Project Implicit data

Figure 2 and Figure 3 show the distribution of the D-Score by gender for both Science and Career in the two samples: LIBRA Project (scientists) and the Project Implicit (general population). The comparison of the score distributions shows a large displacement towards the right (higher values) for male scores versus female scores for the gender-science IAT (Figure 2a). In fact, male scientists working at top research centers have a mean score of 0.34 while women’s mean score is 0.13 as shown in Table 2 (dif=0.21, p<0.001 Student’s t test). This shift is much less pronounced in the general population (Figure 2b and Table 2) (dif=0.08, p<0.001 Student’s t test). Figure 3a and 3b show the distributions of the IAT score for the gender-career association for men and women across the two datasets (Project LIBRA and Project Implicit). In both cases there is a shift of the distribution of women scores toward the right showing that women association of male and career is stronger than the association among men. The difference in the mean IAT score for gender-career of male versus female is almost identical in both datasets (researchers and general population) (dif=-0.1, p<0.001 Student’s t test).

**Figure 2:**
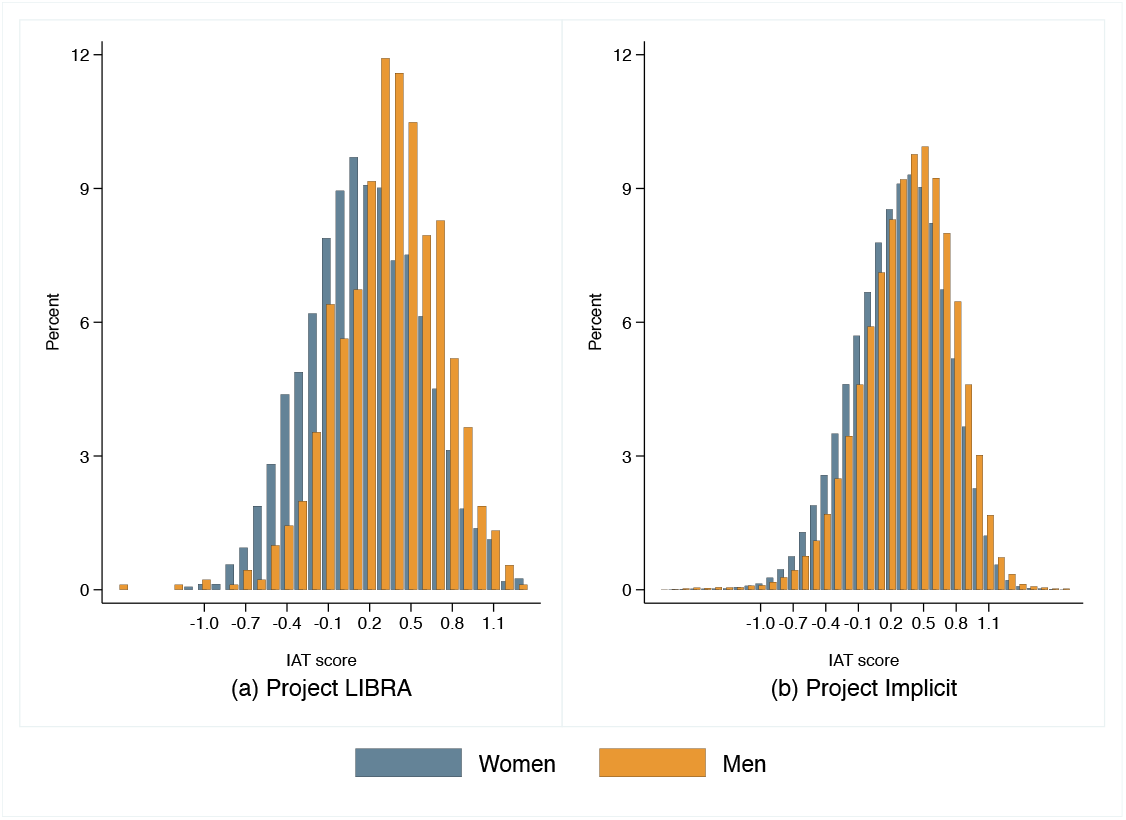
Distribution of the D-score for the gender-science IAT.

**Figure 3:**
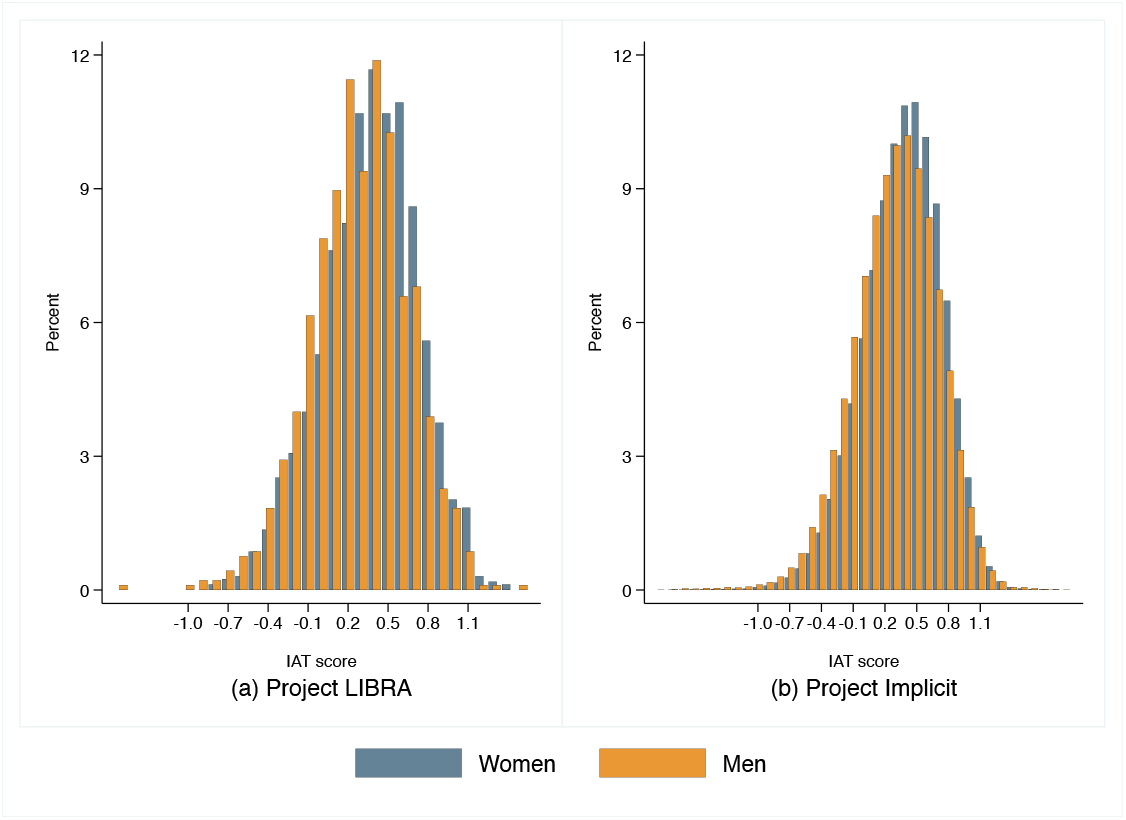
Distribution of the D-score for the gender-career IAT.

## 4 The determinants of the IAT score

In this section we adopt a conditional approach in the analysis of the IAT scores. First of all, we performed a within and between decomposition to understand the basic sources of variation of the IAT (individual or center specific). In order to understand the source of variation in D-scores we run a hierarchical linear model to decompose the total variance into its sources. Equation 4 shows the specification used to obtain the within and between decomposition, and includes two random components, an individual and a research center random component:

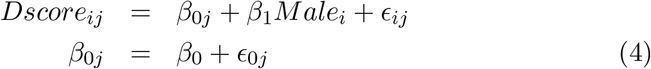

where *Dscore_i_* is the dependent variable for gender-science or gender-career IAT, *β*_0*j*_ is the center specific effect which has two components: a fixed parameter (*β*_0_) and a random component (*ϵ*_0*j*_). The *ϵ_ij_* is the error term. In the second specification of Table 3 we also include the dummy variable *Male_i_*.

**Table 3:** Between/Within center variation of D-Score.

| D-Score | Science |  | Career |  |
| --- | --- | --- | --- | --- |
|  | (1) | (2) | (1) | (2) |
| Male |  | 0.181***<br>(0.017) |  | -0.093***<br>(0.015) |
| Constant | 0.227***<br>(0.013) | 0.161***<br>(0.015) | 0.341***<br>(0.008) | 0.375***<br>(0.010) |
| *Random-effects Parameter* |  |  |  |  |
| Research Center: SD Constant | 0.036 | 0.040 | 0.007 | 0.012 |
| Individual: SD Residual | 0.401 | 0.391 | 0.364 | 0.362 |
| Observations | 2,394 | 2,387 | 2,430 | 2,422 |

Standard errors in parentheses. P-values are exact values calculated for a two sided test of the parameter being equal to 0. \* $p < 0.05$ , \*\* $p < 0.01$ , \*\*\* $p < 0.001$ .
Source: Our elaboration on Project LIBRA data

Using a results of Table 3 we can conclude that most of the variation comes from the individual component. The hierarchical model with two random components shows that the research center effect explains less than 1% of the variance of the gender-science IAT D-score. The explanatory power of the centers’ identity on the gender-career IAT D-score is even smaller than in the gender-science test. In general top research centers attract researchers from many different countries and, therefore, we should not expect to see cultural differences reflected in the IAT score related with the center in which scientists work.

Since almost all the variability of the implicit association between gender and science, and gender and career, comes from individual differences, it is interesting to analyze the determinants at the individual level. Table 4 reports the estimation of the determinants of the gender-science score for the sample of non-administrative staff^6^. The first column includes the basic specification. As we already noticed using descriptive statistics, age increases the implicit association of male and science. The coefficient is statistically significant at the usual level of confidence (*β*=0.004, P<0.01 Student’s t test): a difference of 10 years in the age of a researcher increases the IAT D-score by 0.04 points, or 16% of the unconditional mean. The most relevant variable is, as expected, gender. Conditional on the rest of the variables, male researchers have a male-science IAT D-score above half the unconditional average (*β*=0.20, P<0.001 Student’s t test). The post-doc indicator has a negative coefficient showing that individuals in this group have less gender-science implicit association (*β*=-0.06, P<0.01 Student’s t test). The results are basically unaffected by the inclusion of a dummy variable to control for the research center (column 2 of Table 4).

**Table 4:**
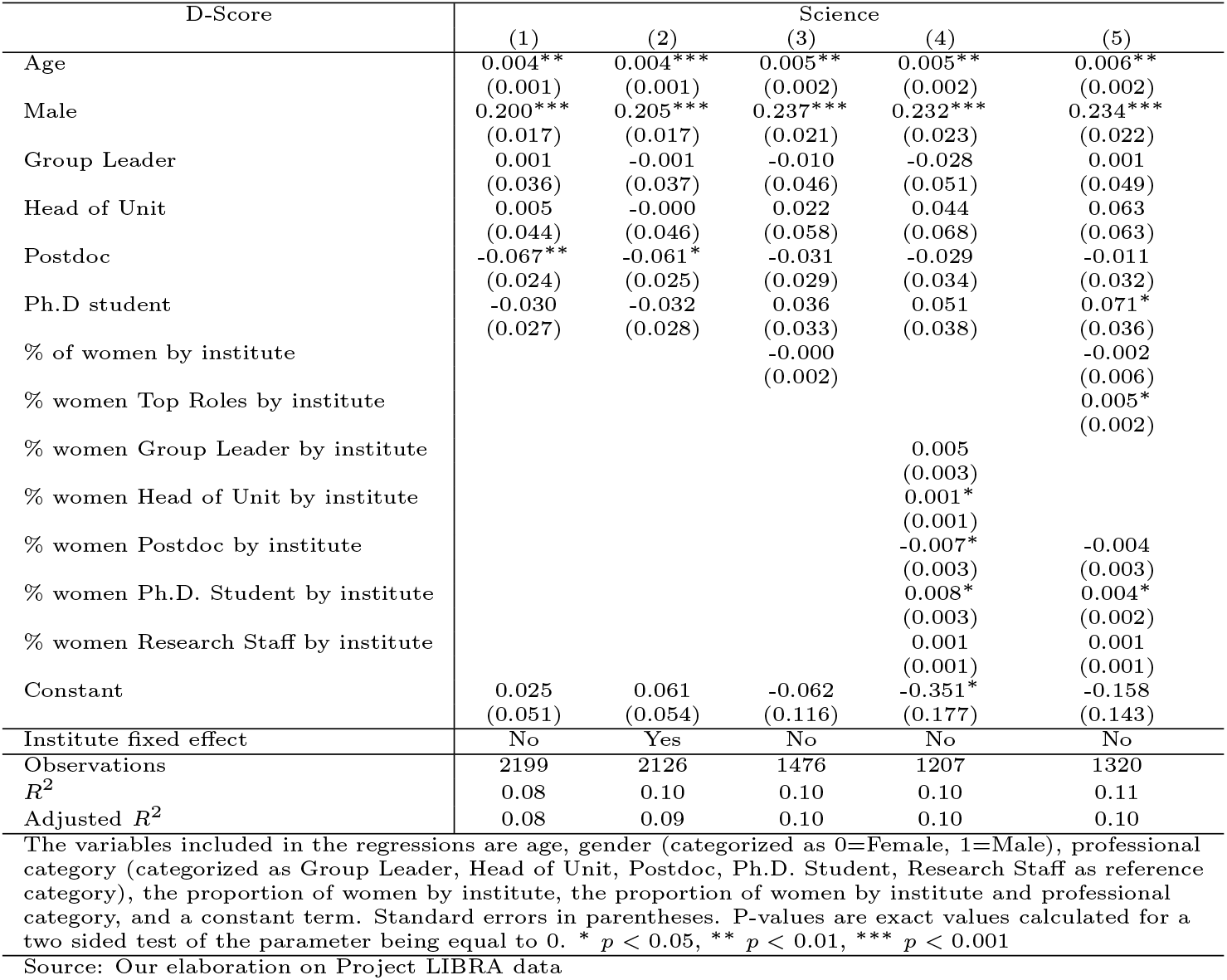
Determinants of D-Score: Science. LIBRA non-administrative staff.

One potentially important explanation of the IAT score are interacting sociocultural factors. Previous research on national differences in gender-science stereotypes shows that a higher female employment in the research sector in a particular country reduces the explicit national gender-science stereotype but does not affect the implicit stereotype (Miller et al., 2015). Even countries located at the top of the ranking of overall gender equity present strong gender-science implicit stereotypes if the field is dominated by men. We can control for this type of effect at the center level by adding the proportion of women in different positions (Column 3 of Table 4). The proportion of female employment in a particular center does not have any significant influence on individual IAT scores. To investigate further the role of women at the level of research centers, Columns 4 and 5 of Table 4 consider separately the proportion of women by professional category. Column 4 of Table 4 shows that the proportions of women that are head of units (*β*=0.001, P<0.05 Student’s t test) have a positive and significant impact on the male-science association at the individual level, though small. This effect is confirmed by the aggregation of both groups as top roles in Column 5 of Table 4. The post-doc effect found in the specifications of Columns 1 and 2 loses statistical significance when center-specific variables are included in the regression (Columns 3 to 5 of Table 3). Instead, the proportion of women among the postdoc group decreases the male-science association, although the effect is not very intense (*β*=-0.007, P<0.05 Student’s t test). Including these additional covariates increases the size of the effect of gender on the gender-science IAT D-score.

The determinants of the gender-career IAT score among researchers are considered in Table 5^7^. In this case males have a lower tendency to associate male and scientific career than female (*β*=-0.08, P<0.001 Student’s t test). This result is very consistent across different specifications. The postdoc and Ph.D. students have also a lower gender-career association than any other positions in the research centers. The effect is large and statistically significant. Considering all the covariates (column 5) the impact of being a postdoc or a Ph.D. student is, in absolute value, even larger than the effect of male on the gender-career IAT (for postdoc *β*=-0.097, P<0.001 Student’s t test; for Ph.D. students *β*=-0.095, P<0.01 Student’s t test). In the case of the gender-career IAT score the administrative staff does not have any significant difference with respect to the rest of the employees. The percentage of women among employees increases the gender-career association (*β*=0.01, P<0.05 Student’s t test) although the effect is small.

**Table 5:** Determinants of D-Score: Career.LIBRA non-administrative staff.

| D-Score | Career |  |  |  |  |
| --- | --- | --- | --- | --- | --- |
|  | (1) | (2) | (3) | (4) | (5) |
| Age | 0.001<br>(0.001) | 0.001<br>(0.001) | 0.002<br>(0.002) | 0.001<br>(0.002) | 0.001<br>(0.002) |
| Male | -0.089***<br>(0.016) | -0.089***<br>(0.016) | -0.090***<br>(0.020) | -0.094***<br>(0.021) | -0.088***<br>(0.021) |
| Group Leader | -0.045<br>(0.032) | -0.045<br>(0.034) | -0.082<br>(0.043) | -0.058<br>(0.047) | -0.073<br>(0.046) |
| Head of Unit | 0.041<br>(0.040) | 0.053<br>(0.042) | -0.006<br>(0.054) | 0.065<br>(0.063) | 0.039<br>(0.059) |
| Postdoc | -0.051*<br>(0.022) | -0.067**<br>(0.023) | -0.072*<br>(0.028) | -0.072*<br>(0.032) | -0.097**<br>(0.031) |
| Ph.D student | -0.045<br>(0.025) | -0.058*<br>(0.026) | -0.071*<br>(0.032) | -0.075*<br>(0.036) | -0.095**<br>(0.035) |
| % of women by institute |  |  | 0.002<br>(0.002) |  | 0.013*<br>(0.006) |
| % women Top Roles by institute |  |  |  |  | -0.004<br>(0.002) |
| % women Group Leader by institute |  |  |  | 0.000<br>(0.002) |  |
| % women Head of Unit by institute |  |  |  | -0.000<br>(0.001) |  |
| % women Postdoc by institute |  |  |  | -0.002<br>(0.003) | -0.002<br>(0.003) |
| % women Ph.D. Student by institute |  |  |  | 0.004<br>(0.003) | -0.003<br>(0.002) |
| % women Research Staff by institute |  |  |  | 0.001<br>(0.001) | -0.002<br>(0.001) |
| Constant | 0.371***<br>(0.047) | 0.368***<br>(0.050) | 0.242*<br>(0.111) | 0.205<br>(0.165) | 0.183<br>(0.139) |
| Institute fixed effect | No | Yes | No | No | No |
| Observations | 2,246 | 2,163 | 1,474 | 1,216 | 1,326 |
| $R^2$ | 0.02 | 0.03 | 0.03 | 0.03 | 0.03 |
| Adjusted $R^2$ | 0.02 | 0.02 | 0.02 | 0.02 | 0.02 |

The variables included in the regressions are age, gender (categorized as 0=Female, 1=Male), professional category (categorized as Group Leader, Head of Unit, Postdoc, Ph.D. Student, Research Staff as reference category), the proportion of women by institute, the proportion of women by institute and professional category, and a constant term. Standard errors in parentheses. P-values are exact values calculated for a two sided test of the parameter being equal to 0. \* $p < 0.05$ , \*\* $p < 0.01$ , \*\*\* $p < 0.001$
Source: Our elaboration on Project LIBRA data

To complete the analysis of the individual specific determinants of the IAT score among biomedical researchers versus the general population we consider a comparison of their determinants. We include three covariates that are common to both datasets: age, gender, and education. Figure 4 shows the point estimate and the 95% confidence interval of the parameter estimates for each sample. The results are basically identical if we use the subsample of the general population of Project Implicit that includes only people 26 years old, or older, who are Europeans^8^.

**Figure 4:**
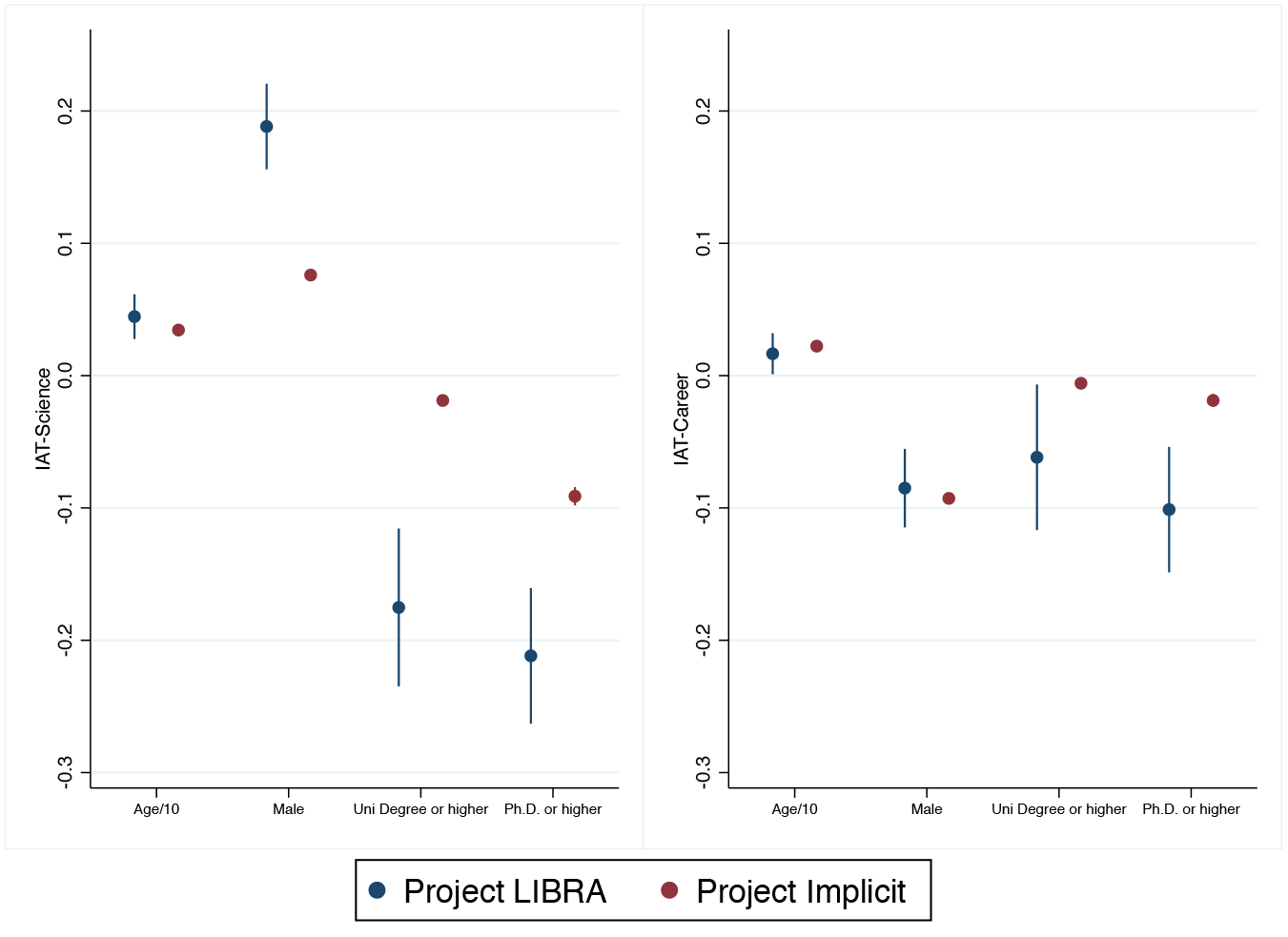
Comparison of the effect of the determinants of the IAT D-Score.

First we analyzed the gender-science IAT score. As we already mentioned in the descriptive section, age has a statistically positive influence on the association gender-science among researchers. This is also the case for the general population although the different between both parameters is not statistically significant. The variable ‘male’ has a much higher coefficient for the sample of researchers (*β*=0.19, P<0.001 Student’s t test) than for the sample of the general population (*β*=0.076, P<0.001 Student’s t test). This implies that male researchers have a much higher association of men and science, relative to female, than the general population. This is consistent with the previous descriptive results that showed that women researchers are the basic reason of the smaller gender-science IAT score in the sample of researchers than in the general population. As expected, the coefficients for age and gender are very similar to the ones obtained in Table 3 and 4, which confirms that working with the common available covariates in both samples does not have a significant impact on the results. Having a university degree or a Ph.D. reduces the male-science association of researchers more than in the case of the general population. In fact, there is a large difference in the effect of a Ph.D. versus a university degree among the general population while the difference is much smaller in the sample of researchers.

The analysis of the comparative determinants of the IAT gender-career score between researchers and general population presented in Figure 4 is also interesting. Age and gender have similar influence on the IAT score across both samples. As we already showed, women tend to depict a higher intensity than men in the association of men and career. The negative effect of a university degree or a Ph.D. on the IAT score is more intense in the case of the sample of researchers than in the general population. Basically, the influence of the different characteristics on the gender-career IAT is very similar in both samples.

## 5 Discussion

Are biomedical researchers less gender biased than the general population? The methodological approach followed by scientists in their research may support their belief that scientific careers are determined by strictly meritocratic procedures. However, implicit biases can hinder the workings of these procedures. Our study provides for the first time an analysis of gender implicit association for science and career among active scientists working in top biomedical research centers across Europe. We show that implicit male-science stereotypes (as measured by the Implicit Association Test) are less prevalent among active scientists working in biomedical research centers than in the general population. However, this result is driven by the low association among female scientists. Male scientists have the same level of implicit male-science association as the general population. Since men dominate decision-making positions, this implicit bias can produce gender inertia at the high levels of the scientific career.

We also found that scientists in top positions have a much higher implicit association for male and science than the general population and that age is a clear determinant. Although the proportion of women in the centers has not impact on individual D-scores, postdoc and PhD students have the lowest scores compared to any other category of employees. Concerning gender-career association, female scientists have a higher score than their fellow male scientists but slightly lower than the general population.

Previous research, using self-reported answers to surveys of academics, argued that to increase women representation among researchers there is need to highlight the importance of effort for pursuing a successful scientific career, instead of innate intellectual gifts (Leslie, Cimpian, Meyer, & Freeland, 2015). Other proposals include to weaken stereotypes highlighting examples of female scientists as part of regular classroom instruction (Miller et al., 2015). The implicit bias among male researchers working in research centers, even stronger in those at top positions, seems to indicate that these strategies would not be enough to revert the gender stereotype in scientific research. In fact previous studies indicated that male scientists are, in general, quite reluctant to accept the existence of a gender bias, what has been described as the myth of “other people are biased not me” (Kang & Kaplan, 2019). Our research shows the existence of implicit gender bias among male scientists, and women in top roles. Therefore, guidelines that enforce a high proportion of women in hiring and promotion committees at the highest levels may not be effective.

Another proposal to address the issue of implicit association of science and male is controlling individual bias directly using diversity training. This strategy seems to be consistent with recent findings (Régner et al., 2019)(Girod et al., 2016). In addition, recent research shows that women are being evaluated less favorably as principal investigators than for the quality of the proposals (Witterman, Hendricks, Straus, & Tannenbaum, 2019). Since selection processes are mostly based on the merits of the candidates, we propose to facilitate a transparent and equitable hiring by asking committees to write detailed discussions on the explicit merits of the candidates for a job/promotion to be reviewed afterwards, in order to limit the scope for unconscious reasons in the selection/promotion process. Most of the time these decisions are taken in oral discussions and, therefore, it is difficult to avoid the influence of implicit biases based on applying different arguments to different candidates depending on their gender. The written documents of the discussions should help to produce consistent decisions over time, and cumulative guidelines for gender neutral decisions that could be analyzed by external experts and improved over time.

## Appendix A The Implicit Association Test

The gender-science IAT measures the relative speed at which the experimental subject can associate categories, such as men or women, and attributes, related with science and liberal arts. In the case of the gender-career IAT the attributes are related with family and career. The relative speed at which the subject can pair categories and attributes, and the number of classification errors, measures implicitly the strength of the association between categories and attributes (see Methods and Materials for details). The response time differences provide a score that can be interpreted as evidence of implicit gender-science, or gender-career, stereotypes. For instance, fast sorting of stereotype-congruent categories and attributes relative to incongruous ones points out to higher association of science and male than science and female.

The scientists in the research centers in our sample performed a computer-based set of trials that allow the measurement of the strength of automatic associations between concepts by analyzing latency in the responses provided to each of the tasks. For analyzing the automatic association of gender with science in comparison to liberal arts, attributes needed to be sorted into categories defined by one or two of the following four concepts *Female*, *Male*, *Science*, and *Liberal arts*. The displayed attributes per concept were as follows: *Female*: Mother, Wife, Aunt, Women, Girl, Female, Grandma, Daughter; *Male*: Man, Son, Father, Boy, Uncle, Grandpa, Husband, Male; *Science*: Astronomy, Math, Chemistry, Physics, Biology, Geology, Engineering; *Liberal arts*: History, Arts, Humanities, English, Philosophy, Music, Literature. The categories were defined differently in seven individual tasks and respondents were asked to sort above mentioned attributes into the two categories as quickly as possible by using the two keys an *E* key for one and *I* for the other category. Items appeared on the screen one at a time and each item belongs to only one category. If the respondent did a mistake, a red X appeared on the screen and the respondent had to press the other key to correct the answer. The seven tasks were as follows: (1) Put a left finger on the *E* key for items that belong to the category *Liberal Arts*. Put the right finger on the *I* key for the items that belong to the category *Science*.; (2) Put a left finger on the *E* key for items that belong to the category *Male*. Put the right finger on the *I* key for the items that belong to the category *Female*.; (3) Use the *E* key for *Liberal Arts* and for *Male*. Use the *I* key for *Science* and for *Female*.; (4) This is the same as the previous part. Use the *E* key for *Liberal Arts* and for *Male*. Use the *I* key for *Science* and for *Female*.; (5) Watch out, the labels have changed position! Use the left finger on the *E* key for *Science*. Use the right finger on the *I* key for *Liberal Arts*.; (6) Use the *E* key for *Science* and for *Male*. Use the *I* key for the *Liberal Arts* and for *Female*.; (7) This is the same as the previous part. Use the *E* key for *Science* and for *Male*. Use the *I* key for the *Liberal Arts* and for *Female*. In the two task blocks 3/4 and 6/7 respondents repeated the same task with the same keys for the same gender but with exchanged second concept. For example, in block 3/4 one category combines *Male* with *Liberal Arts*, while in block 6/7 *Male* is combined with *Science*, and in both blocks items need to be sorted with the key *E*. The difference in average latency between those two task blocks is at the base of the IAT measure. As an example, a faster performance in tasks 6/7 shows a stronger association of *Male* than *Female* with *Science*. The order of the blocks can influence the overall results slightly. If the first block is pairing *Science* and *Male* and then pairing *Science* and *Female*, the results might show a slightly stronger association of *Male* with *Science* than in the reversed order of pairing. Therefore, the IAT foresees two measures for minimizing the order effect. First, in this case giving more practice trials before the second pairing than for the first pairing, and second, the participants are randomly assigned to a specific order. Half of test-takers complete first the *Science* and *Male* block, and then the *Science* and *Female* block, while the other half of test-takers get the opposite order.

For analyzing the automatic association of gender with career in comparison to family the same principles of the IAT apply. While the gender concepts *Male* and *Female* are also used in this test, the concepts *Science* and *Liberal Arts* were replaced by *Career* and *Family*. Related attributes which needed to be sorted were as follows: *Female*: Rebecca, Michelle, Emily, Julia, Anna; *Male*:Ben, Paul, Daniel, John, Jeffrey; *Career*: Career, Corporation, Salary, Office, Professional, Management, Business; *Family*: Wedding, Marriage, Parents, Relatives, Family, Home, Children. The seven tasks were as follows: (1) Put a left finger on the *E* key for items that belong to the category *Family*. Put the right finger on the *I* key for the items that belong to the category *Career*. (2) Put a left finger on the *E* key for items that belong to the category *Male*. Put the right finger on the *I* key for the items that belong to the category *Female*. (3) Use the *E* key for *Family* and for *Male*. Use the *I* key for *Career* and for *Female*. (4) This is the same as the previous part. Use the *E* key for *Family* and for *Male*. Use the *I* key for *Career* and for *Female*. (5) Watch out, the labels have changed position! Use the left finger on the *E* key for *Career*. Use the right finger on the *I* key for *Family*. (6) Use the *E* key for *Career* and for *Male*. Use the *I* key for the *Family* and for *Female*. (7) This is the same as the previous part. Use the *E* key for *Career* and for *Male*. Use the *I* key for the *Family* and for *Female*.

## Supplementary material

**Table S1:**
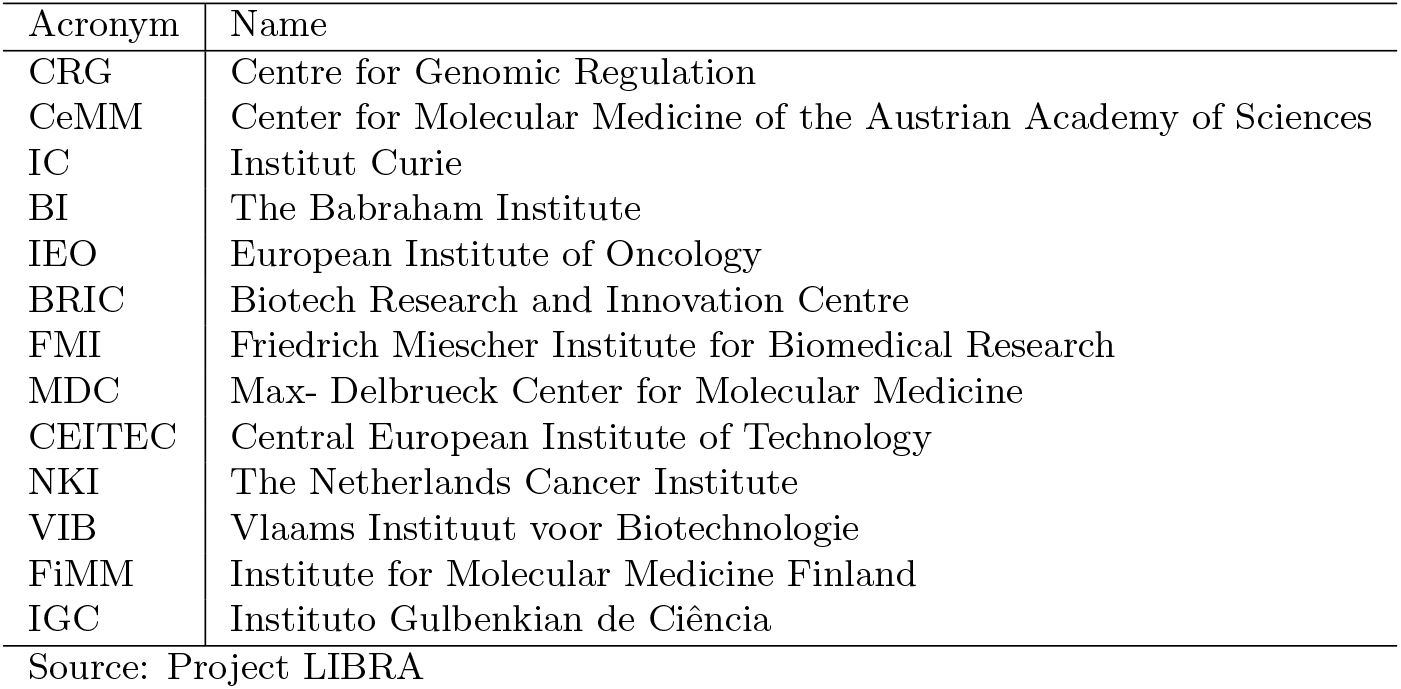
Names and acronyms of the research centers.

It is interesting to check the robustness of the results (Table 2) to the full sample of employees at research centers (including administrative staff). The basic results are robust to including the administrative staff although the difference between the gender-science IAT scores of men and women (dif=0.17, p<0.01 Student’s t test) is a bit smaller when considering all the employees as shown in Table S2.

In fact, the scores for males with or without administrative staff are identical whereas female scientists have an even lower score when considered on their own (without female administrative staff). Altogether female scientists have a significantly lower score for male-science association than their fellow men scientists or other females administrative staff members working in the same institutes.

**Table S2:** Comparison by Gender: full sample.

|  | Project LIBRA |  | Project Implicit |  |
| --- | --- | --- | --- | --- |
|  | Female | Male | Female | Male |
| Mean D-score: Science | 0.17 | 0.34 | 0.28 | 0.36 |
| Mean D-score: Career | 0.38 | 0.28 | 0.38 | 0.29 |

Note: In this table the Project LIBRA sample includes the administrative staff.
Source: Our elaboration on Project LIBRA and Project Implicit data

Restricting the sample even further to analyze only researchers in top roles (Table S3) shows that their implicit association of male and science is higher than in the general population. This substantial change could be related with the higher proportion of men among researchers with top roles, but also with a higher association of male and science among women in top roles versus women in other positions.

**Table S3:**
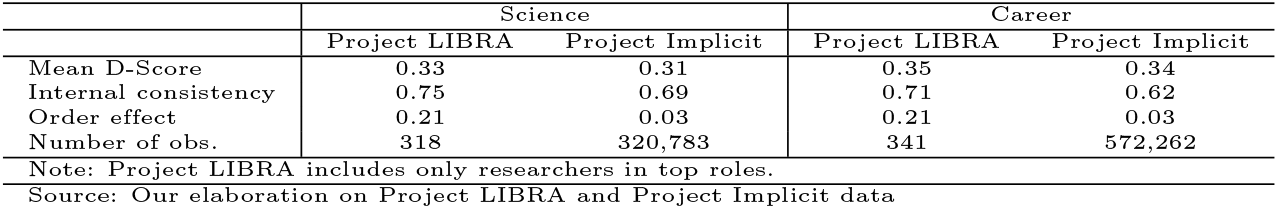
Comparison between researchers in top roles and general population.

Another important determinant is the age of the researchers. We should notice that, since we are working with a cross section of subjects, it is not possible to identify separately the cohort effect from the effect of age. However, this is less of an issue if we want to compare the effect of age between researchers and general population. The correlation of age with the gender-science IAT D-score is higher in the case of researchers than in the general population (Table S4). In contrast, the correlation of age with the gender-career score is identical.

**Table S4:**
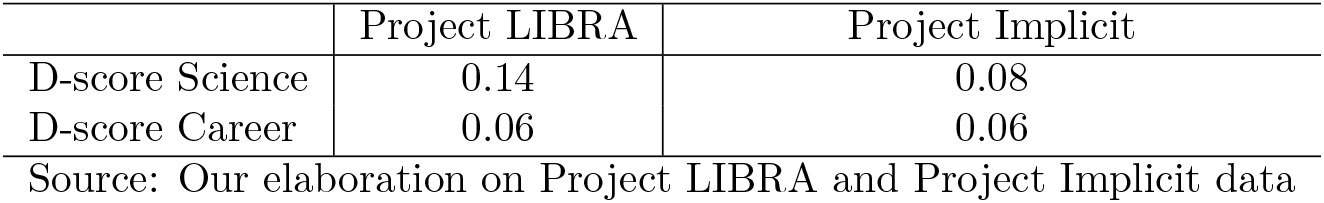
Correlation of the IAT D-Score with age.

|  | Project LIBRA | Project Implicit |
| --- | --- | --- |
| D-score Science | 0.14 | 0.08 |
| D-score Career | 0.06 | 0.06 |

Source: Our elaboration on Project LIBRA and Project Implicit data

The basic results are also robust using the full sample, which includes the administrative staff of the centers (Table S5 and Table S6). Table S5 shows that males working at the research centers have a higher D-score than females being the estimates (*β*=0.18 to *β*=0.23 depending on the specification, P<0.001 Student’s t test) quite similar to the ones presented in Table 6, which do not included the administrative staff in the sample. The personnel in the administrative group shows a higher level of association of male and science than researchers (*β*=0.17 to *β*=0.24, P<0.001 Student’s t test), despite the fact that women represent the largest proportion in this group. However, Table S6 shows that the administrative staff do not have a robust differential effect on the gender-career IAT D-score. Finally the effect of gender on the score is not affected by the inclusion of the administrative staff in the sample once all the conditioning variables are included in the specification.

**Table S5:** Determinants of the D-Score: Science. LIBRA full sample.

| D-Score | Science |  |  |  |  |
| --- | --- | --- | --- | --- | --- |
|  | (1) | (2) | (3) | (4) | (5) |
| Age | 0.004**<br>(0.001) | 0.004***<br>(0.001) | 0.005**<br>(0.001) | 0.005**<br>(0.002) | 0.006**<br>(0.002) |
| Male | 0.186***<br>(0.017) | 0.193***<br>(0.017) | 0.233***<br>(0.020) | 0.216***<br>(0.024) | 0.220***<br>(0.023) |
| Group Leader | 0.008<br>(0.035) | 0.006<br>(0.036) | -0.003<br>(0.045) | -0.007<br>(0.054) | 0.017<br>(0.052) |
| Head of Unit | 0.012<br>(0.043) | 0.008<br>(0.045) | 0.029<br>(0.057) | 0.073<br>(0.072) | 0.079<br>(0.066) |
| Postdoc | -0.066**<br>(0.024) | -0.059*<br>(0.025) | -0.032<br>(0.029) | -0.002<br>(0.037) | 0.008<br>(0.034) |
| Ph.D student | -0.032<br>(0.026) | -0.033<br>(0.027) | 0.031<br>(0.032) | 0.072<br>(0.041) | 0.083*<br>(0.039) |
| Administrative | 0.178***<br>(0.031) | 0.153***<br>(0.032) | 0.203***<br>(0.040) | 0.217***<br>(0.047) | 0.236***<br>(0.044) |
| % of women by institute |  |  | -0.000<br>(0.002) |  | 0.025<br>(0.016) |
| % women Top Roles by institute |  |  |  |  | 0.000<br>(0.004) |
| % women Group Leader by institute |  |  |  | 0.005*<br>(0.002) |  |
| % women Head of Unit by institute |  |  |  | 0.002**<br>(0.001) |  |
| % women Postdoc by institute |  |  |  | -0.008*<br>(0.003) | -0.014**<br>(0.005) |
| % women Ph.D. Student by institute |  |  |  | 0.008*<br>(0.004) | -0.003<br>(0.005) |
| % women Research Staff by institute |  |  |  | 0.001<br>(0.001) | -0.004<br>(0.003) |
| % women Administrative by institute |  |  |  | 0.000<br>(0.001) | 0.003<br>(0.003) |
| Constant | 0.042<br>(0.046) | 0.069<br>(0.050) | -0.045<br>(0.109) | -0.404*<br>(0.172) | -0.664*<br>(0.320) |
| Institute fixed effect | No | Yes | No | No | No |
| Observations | 2,473 | 2,372 | 1,627 | 1,151 | 1,275 |
| $R^2$ | 0.09 | 0.10 | 0.11 | 0.11 | 0.11 |
| Adjusted $R^2$ | 0.08 | 0.09 | 0.10 | 0.10 | 0.10 |

The variables included in the regressions are age, gender (categorized as 0=Female, 1=Male), professional category (categorized as Group Leader, Head of Unit, Postdoc, Ph.D. Student, Administrative, Research Staff as reference category), the proportion of women by institute, the proportion of women by institute and professional category, and a constant term. Standard errors in parentheses. P-values are exact values calculated for a two sided test of the parameter being equal to 0. \* $p < 0.05$ , \*\* $p < 0.01$ , \*\*\* $p < 0.001$
Source: Our elaboration on Project LIBRA data

**Table S6:** Determinants of the D-Score: Career. LIBRA full sample.

| D-Score | Career |  |  |  |  |
| --- | --- | --- | --- | --- | --- |
|  | (1) | (2) | (3) | (4) | (5) |
| Age | 0.001<br>(0.001) | 0.001<br>(0.001) | 0.002<br>(0.001) | 0.002<br>(0.002) | 0.002<br>(0.002) |
| Male | -0.084***<br>(0.015) | -0.085***<br>(0.016) | -0.084***<br>(0.019) | -0.079***<br>(0.022) | -0.076***<br>(0.022) |
| Group Leader | -0.049<br>(0.032) | -0.049<br>(0.033) | -0.085*<br>(0.042) | -0.086<br>(0.050) | -0.096*<br>(0.049) |
| Head of Unit | 0.037<br>(0.039) | 0.046<br>(0.041) | -0.009<br>(0.053) | 0.006<br>(0.065) | -0.017<br>(0.062) |
| Postdoc | -0.051*<br>(0.022) | -0.068**<br>(0.023) | -0.072**<br>(0.028) | -0.092**<br>(0.035) | -0.112***<br>(0.033) |
| Ph.D student | -0.043<br>(0.024) | -0.057*<br>(0.025) | -0.071*<br>(0.031) | -0.094*<br>(0.039) | -0.115**<br>(0.038) |
| Administrative | 0.064*<br>(0.028) | 0.052<br>(0.030) | 0.048<br>(0.038) | 0.043<br>(0.044) | 0.027<br>(0.042) |
| % of women by institute |  |  | 0.003<br>(0.002) |  | 0.032*<br>(0.015) |
| % women Top Roles by institute |  |  |  |  | -0.009*<br>(0.004) |
| % women Group Leader by institute |  |  |  | 0.001<br>(0.002) |  |
| % women Head of Unit by institute |  |  |  | -0.000<br>(0.001) |  |
| % women Postdoc by institute |  |  |  | -0.003<br>(0.003) | -0.007<br>(0.005) |
| % women Ph.D. Student by institute |  |  |  | 0.008*<br>(0.004) | -0.007<br>(0.005) |
| % women Research Staff by institute |  |  |  | 0.002<br>(0.001) | -0.006<br>(0.003) |
| % women Administrative by institute |  |  |  | -0.002<br>(0.001) | 0.003<br>(0.003) |
| Constant | 0.359***<br>(0.043) | 0.358***<br>(0.046) | 0.223*<br>(0.104) | 0.126<br>(0.161) | -0.254<br>(0.308) |
| Institute fixed effect | No | Yes | No | No | No |
| Observations | 2,520 | 2,407 | 1,624 | 1,147 | 1,267 |
| R<sup>2</sup> | 0.03 | 0.04 | 0.04 | 0.05 | 0.05 |
| Adjusted R<sup>2</sup> | 0.03 | 0.03 | 0.03 | 0.04 | 0.04 |

The variables included in the regressions are age, gender (categorized as 0=Female, 1=Male), professional category (categorized as Group Leader, Head of Unit, Postdoc, Ph.D. Student, Administrative, Research Staff as reference category), the proportion of women by institute, the proportion of women by institute and professional category, and a constant term. Standard errors in parentheses. P-values are exact values calculated for a two sided test of the parameter being equal to 0. \* $p < 0.05$ , \*\* $p < 0.01$ , \*\*\* $p < 0.001$
Source: Our elaboration on Project LIBRA data

The previous analysis uses age as a covariate. We also analyzed this effect using age grouped by cohort. Figure S1 depicts the coefficients of each cohort, and its corresponding standard deviation, coming from a regression that includes all the covariates of column 5 of Table 3 and 4 but substitute age by age-cohorts.

**Figure S1:**
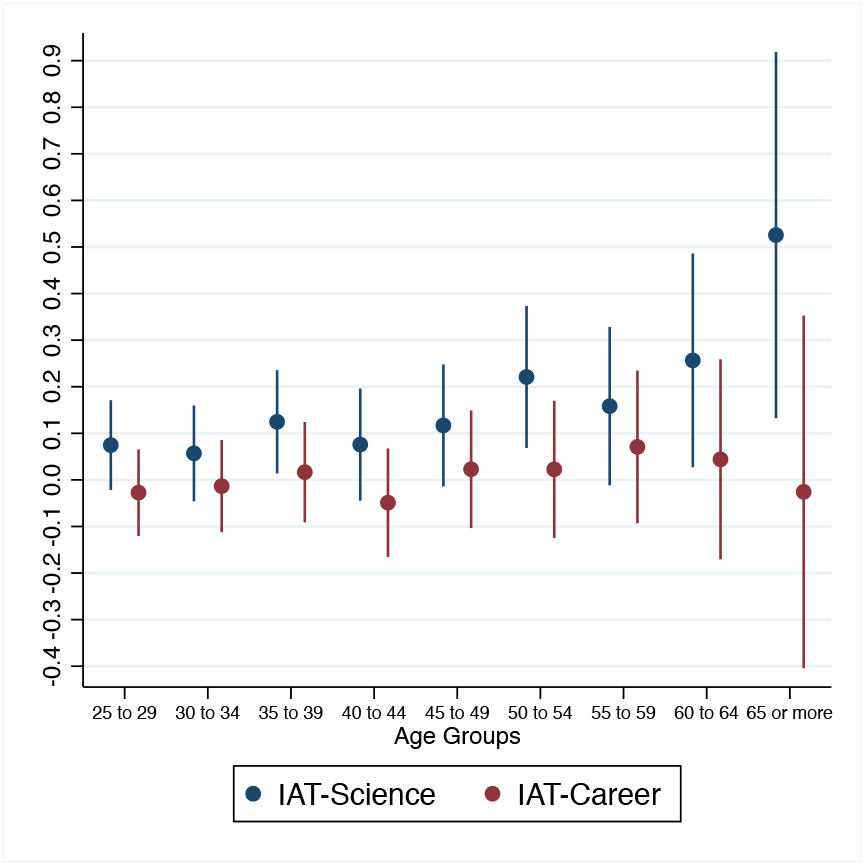
Cohort effect on the IAT LIBRA.

The last two columns of Table S7 include the analysis of the IAT scores for a subsample of the general population of Project Implicit that includes only people 26 years old, or older, who are Europeans. Since the scientists analyzed in this paper work in European research centers, this sub-sample is the closest, in the Project Implicit data, to the researchers who took our IAT.^1^ With respect to the difference in gender-science score between male and females the results are quite similar to the ones obtained using all the respondents of the Project Implicit: the difference is around 0.19 among the scientists while it is 0.08 for the whole population of the Project Implicit, and 0.09 for the sub-group of European citizens 26 years old or older. The effect of age on the score is similar to the one obtained in the sample of researchers. However, the European male graduates have a much lower (−0.05) difference with respect to females in the gender-science IAT score than the difference obtained in the full sample of the Project Implicit data (−0.09) or in the sample of researchers (−0.21). The gender-career IAT shows similar results in all the groups with respect to the effect of gender: women have a higher intensity of association of male and career than men in the tree groups (scientists, general population and Europeans).

**Table S7:**
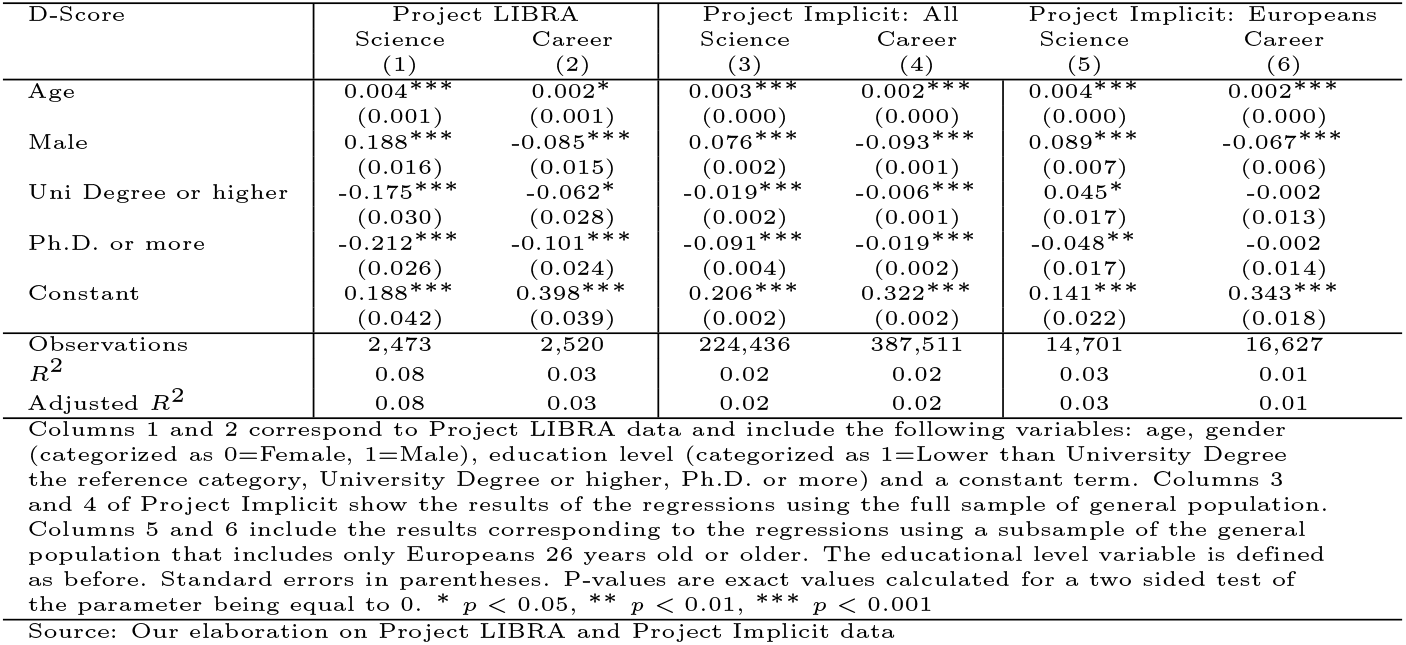
Determinants of D-Score: Project LIBRA and Project Implicit.

## Footnotes

1 Appendix A presents a detailed description of the test and its application.

2 Supplementary Material, Table S1.

3 The Supplementary Material shows the robustness of these results to the whole sample of workers at the research centers.

4 Table S2 in the Supplementary Material shows that the results of Table 2 are robust to including all the workers and not only the researchers at the center.

5 Table S3 in the Supplementary Material shows that, in the sample of researchers in top roles (group leaders and head of units) the difference in the IAT of science between scientist and the general population is not statistically significant. This is mostly due to the fact that researcher in top roles are predominantly men.

6 Table S5 shows the results of the estimation including all the employees of the research centers.

7 Table S6 shows the results of the estimation including all the employees of the research centers.

8 Table S7 in the Supplementary Material presents the estimations. The last two columns of Table S7 include the results considering among the general population only Europeans older than 26 years old.

1 Notice that there are also US citizen working at these research centers.

